# Single session of rTMS enhances brain metastability and intrinsic ignition

**DOI:** 10.1101/2022.08.14.503887

**Authors:** Sujas Bhardwaj, Rajanikant Panda, Rose Dawn Bharath, Albert Stezin, Sunil Kumar Khokhar, Shweta Prasad, Vidhi Tyagi, Nitish Kamble, Keshav Kumar, Netravathi M, Ravi Yadav, Rajan Kashyap, Kaviraja Udupa, Jitka Annen, Steven Laureys, Gustavo Deco, Pramod Kumar Pal

**Affiliations:** Neuroimaging and Interventional Radiology, National Institute of Mental Health and Neurosciences (NIMHANS), Bengaluru, India; Neurology, National Institute of Mental Health and Neurosciences (NIMHANS), Bengaluru, India; Music Cognition Lab, Neurobiology Research Centre, National Institute of Mental Health and Neurosciences (NIMHANS), Bengaluru, India; Coma Science Group, GIGA-Consciousness, University of Liège, Belgium; Centre du Cerveau^2^, University Hospital of Liège, Belgium; Clinical Neurosciences, National Institute of Mental Health and Neurosciences (NIMHANS), Bengaluru, India; Centre for Brain Research, Indian Institute of Science, Bengaluru, India; Clinical Psychology, National Institute of Mental Health and Neurosciences (NIMHANS), Bengaluru, India; Neurophysiology, National Institute of Mental Health and Neurosciences (NIMHANS), Bengaluru, India; Center for Brain and Cognition, Computational Neuroscience Group, Department of Information and Communication Technologies, Universitat Pompeu Fabra, Barcelona, 08018, Spain

**Author notes:** **Corresponding Author:** Dr. Rose Dawn Bharath, Professor, Neuroimaging and Interventional Radiology, NIMHANS, Bengaluru-29, India. [Contact; ]. Equally contributed to the manuscript (Sujas Bhardwaj, Rajanikant Panda, Rose Dawn Bharath).

**Keywords:** repetitive Transcranial Magnetic Stimulation (rTMS), Resting State fMRI, Dynamic Functional Connectivity, Metastability, Intrinsic Ignition

## Abstract

**Background:** Emerging evidence support the view that brain stimulation might improve essential tremor (ET) by altering brain networks and facilitating plasticity. Yet, we are still missing a mechanistic explanation of the whole brain dynamics underlying these plasticity defining changes.

**Method:** In this study, we explored the effect of low-frequency repetitive transcranial magnetic stimulation (rTMS) over left primary motor cortex (L-M1) on functional connectivity dynamics (FCD) in patients with ET. Resting-state fMRI (RsfMRI) was acquired before and after a single session of rTMS in 30 patients with ET and compared with RsfMRI of 20 age and gender matched healthy controls (HCs). We have measured the effect of brain stimulation using network topological re-organization through whole brain integration and segregation, brain stability and capacity of neural propagation through metastability and intrinsic ignition.

**Results:** Patients with ET had altered FCD measures compared to controls. After a single session rTMS, the brain connectivity measures approached normality and patients with ET revealed significantly higher integration, lower segregation with higher metastability and increased intrinsic ignition.

**Conclusion:** Brain metastability and intrinsic ignition measures could be valuable tools in appreciating mechanisms of brain stimulation in ET and other neurological diseases.

## 1. Introduction

Neurological disorders are characterized by several disabling symptoms for which effective, mechanism-based treatments remain elusive and more advanced non-invasive therapeutic methods are being explored. Repetitive transcranial magnetic stimulation (rTMS) is a widely used classical non-invasive brain stimulation (NIBS) method which has been quite useful in management of drug resistant psychiatric disorders such as depression ^1-4^, mood disorders ^5,6^, obsessive compulsive disorder ^7-10^, posttraumatic stress disorder (PTSD) ^11-13^ etc. These studies present mixed results in improvement of patient symptoms or clinical scores. Over the last three decades, TMS has helped us in understanding pathophysiology of many neurological disorders. TMS studies in essential tremor (ET), the most common tremor syndrome, have been helpful in demonstrating cerebello-thalamo-cortical circuitry (CTC) that is likely to be involved in the generation of tremor ^14^. Pathophysiological insights into tremor syndromes have been supportive in selecting appropriate stimulation regions and rTMS parameters in the management of these diseases clinically ^15^.

Resting state fMRI (RsfMRI) methods in duplicate, before and after brain stimulation, is a popular method to assess the neurobiology of brain stimulation owing to its non-invasive nature, capability for whole brain analysis, better spatial resolution of deep-seated brain regions, ease of acquisition, repeatability, and lack of any known adverse effects. Majority of the studies, using RsfMRI reveal a diffuse increased whole brain connectivity immediately after stimulation irrespective of the frequency of stimulation ^16-19^. Though some studies reveal decreased or no changes in connectivity ^20-22^, most of the studies have reported increased connectivity that extended beyond the stimulated region or networks ^17,23,24^. This suggests that the effects of rTMS could either spread through anatomical tracts ^25^ or entrain brain oscillations increasing neural synchrony as a whole ^26^. Stimulation of “task irrelevant” brain area like vertex ^27^ or sham stimulation revealed no changes in connectivity ^28^, reiterating the validity of this tool in measuring changes induced by rTMS. Another interesting point is that the majority of studies report increased connectivity both after a single session of rTMS and after rTMS therapy indicating the potential of using single session rTMS for research. One study ^23^ explored network reorganization and found increased clustering coefficient and reduced path-length suggesting enhanced small-world characteristics of brain network after a single session of LF-rTMS. Recent studies also indicate that these changes could be region specific, as the sensory cortex stimulation reveals different connectivity profile compared to motor cortex ^29^. Homogeneous study groups, individualized target identification, uniform acquisition and analysis methods and longitudinal studies to measure sustainability of these changes are still required to completely understand underlying network reorganization following rTMS.

The concept of brain function being a result of metastable large scale network interactions is being explored using brain connectivity measures. This view suggests that the brain operates in a state of dynamic balance, with different regions constantly communicating and competing with each other in order to perform its functions. One of the key pieces of evidence supporting this concept comes from studies using resting state functional MRI (fMRI). These studies have shown that brain exhibits a number of distinct, highly correlated networks that are active even in the absence of an explicit task, and that these networks are in a constant state of flux, with the strength of their connections changing over time ^30,31^.

Various dynamic functional connectivity measures, such as reproducible patterns of sliding window correlations, single-volume co-activation patterns, and repeating sequence of BOLD activity, have been used to explore the multistability of the brain. Multistability refers to the observation that functional connectivity patterns between brain regions can switch between different states over time. This concept has been used to explain a variety of phenomena in the brain, including the ability of the brain to rapidly switch between different cognitive states in response to changing task demands. Recently, a new concept of, metastability has been put forward to describe the continuously changing and evolving nature of brain connectivity over time. Unlike multistability, which refers to the switching between different states, metastability refers to the idea that functional connectivity patterns are in a state of constant change ^31-35^. Another variable, intrinsic ignition is a measure of interplay of two opposing tendencies in the brain connectivity: the tendency towards coupling and the tendency towards independence. Coupling refers to the tendency of different regions in the brain to form strong functional connections, while independence refers to the tendency of different regions to maintain their own unique patterns of activity. This interplay of opposing tendencies is thought to give rise to the phenomenon of intrinsic ignition, where the brain shifts between multiple stable states in response to changing inputs. Unlike traditional methods, which typically assume the presence of fixed points of equilibrium, intrinsic ignition methods do not rely on this assumption, instead recognizing that the brain’s dynamics are inherently dynamic and adaptable ^36,37^. High intrinsic ignition corresponds to a brain network that is highly adaptable and capable of processing a large amount of information. This is achieved through a rich and flexible brain dynamics, where the network can rapidly switch between multiple stable states and form new connections. This leads to a higher capacity to process event information and respond to changing environmental demands. In contrast, low intrinsic ignition corresponds to a brain network that is less flexible and less adaptable. This is characterized by rigid network interactions, reduced neural communication, and a reduced capacity to process information ^36^.

Though significant RsfMRI imaging-based literature has accrued in the last couple of years revealing modulatory functional connectivity changes after a single session of rTMS ^38^, however, its underlying spatiotemporal alteration using recent measures like metastability and intrinsic ignition is still unclear. In this regard, we employ dynamic functional connectivity assessments to explore whether the stimulatory effect of rTMS on L-M1 in ET can alter metastability, intrinsic ignition and thus the dynamic repertoire of the brain.

## 2. Result

A single session of rTMS was found to increase integration and decrease segregation in patients with ET. Metastability and intrinsic ignition were found to be higher after rTMS at a group level and at individual patient level. Details of these results are as follows:

### 2.1 M1 static connectivity

Before assessing the dynamic functional connectivity measures, we first assesed the time averaged functional connectivity (static) of left primary motor cortex (L-M1) after rTMS using seed to voxel-based connectivity measures. We found increased connectivity (p>0.05 FDR) of left M1 seed with bilateral pre and post central gyrus, central opercula cortex, left temporal pole, left temporal fusiform and right superior middle temporal cortex (Figure 1).

**Figure 1.**
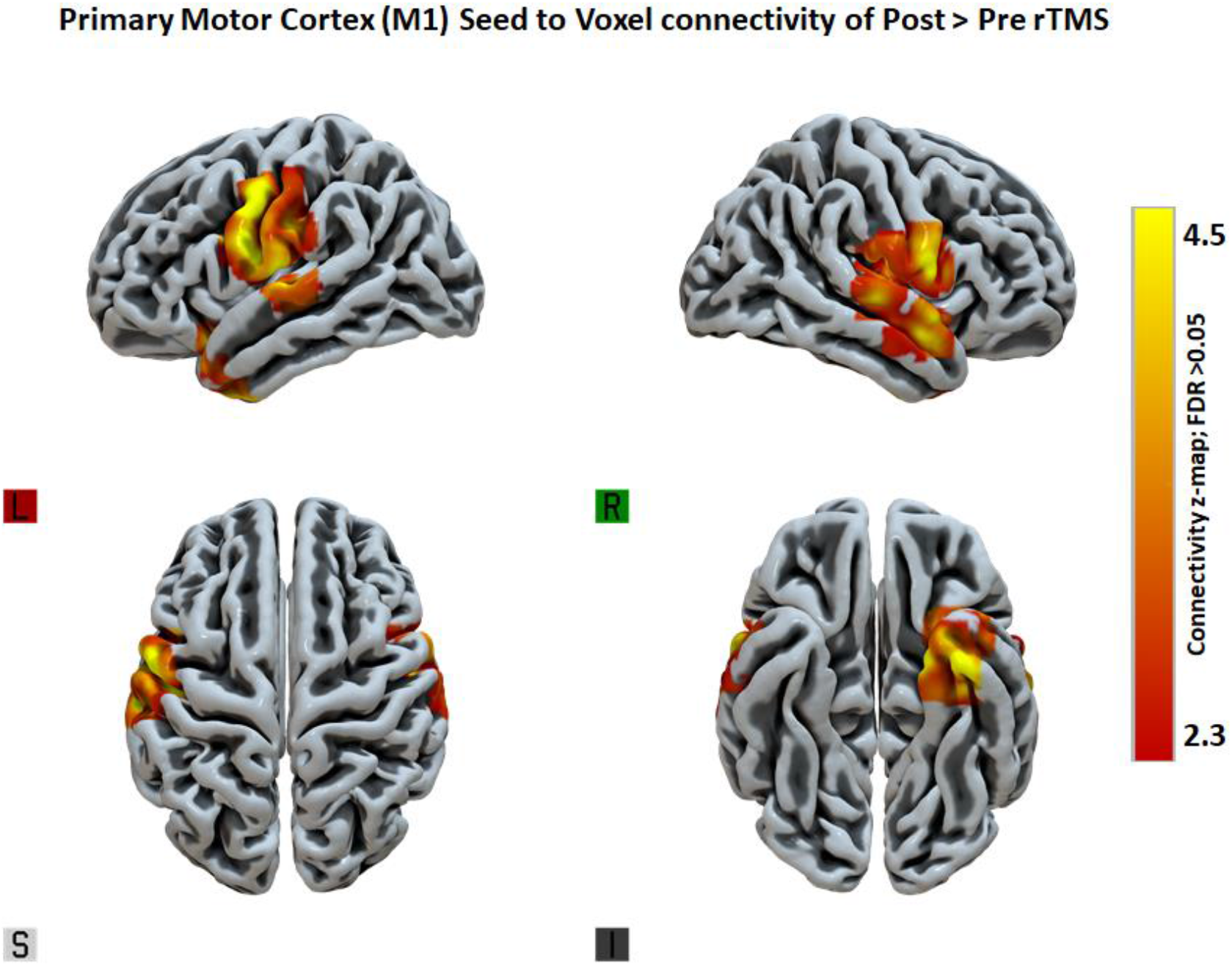
Static functional connectivity of left primary motor cortex (L-M1) for post rTMS compared to pre rTMS in the ET patients.

### 2.2 Dynamic Functional connectivity measure

First to characterize the connectivity differences, we represent the mean phase interaction measures across time for each group/conditions (Figure 2). Followed to this, we quantified recurrence of the phase-interaction patterns through functional connectivity dynamics (FCD). The whole brain FCD was lower in ET (0.24±0.4; p=5.69E-06) than controls (0.39±0.19). After rTMS ET group showed significantly increased mean connectivity (0.48±0.12; p=1.76E-13) (Figure 3.a).

**Figure 2.**
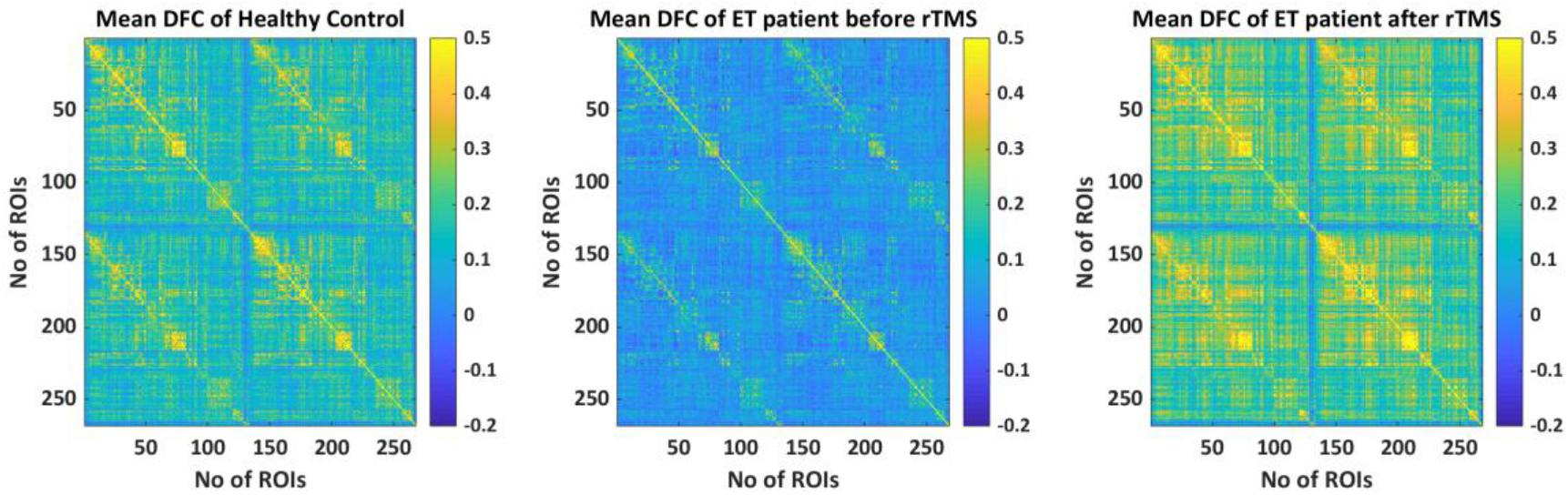
Mean dynamic functional connectivity (DFC) matrix of (a) healthy controls (b) ET before rTMS (c) ET after rTMS stimulus. The mean DFC matrix shows ET having lower functional connectivity as compared to healthy controls, which increased after rTMS.

**Figure 3.**
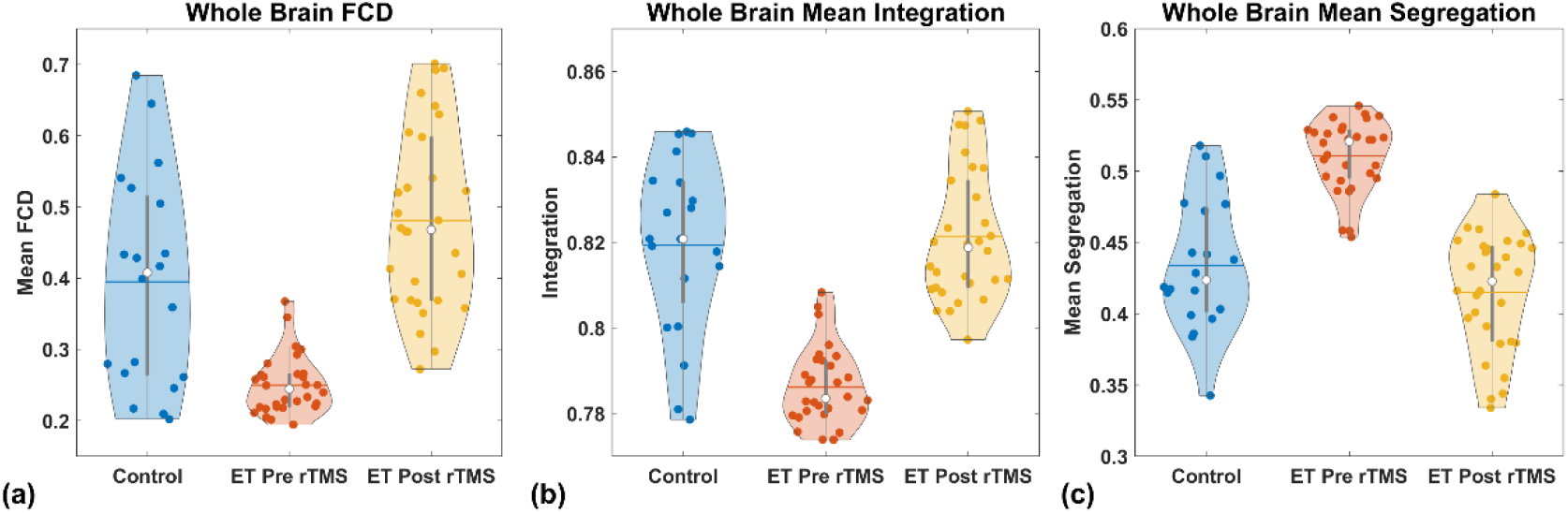
Whole brain (a) FCD (b) network integration and (c) segregation. ET has lower mean FCD, network integration, and higher segregation as compared to healthy controls. After rTMS, mean FCD and brain network integration increased, and segregation decreased in ET.

### 2.3 Whole Brain Integration and Segregation

The mean value of the integration was significantly lower in ET (controls: 0.82±0.02; ET patients in pre rTMS: 0.79±0.009; p= 4.14E-10); after rTMS the brain integration revealed significant increase (0.82±0.01; p=1.419E-14) (Figure 3.b). On the other hand, the brain segregation showed the opposite tendency of integration with increased segregation noted in patients at baseline (controls: 0.43±0.04; ET patients in pre rTMS: 0.51±0.02; p=8.84E-10), and decreased after rTMS (0.41±0.04; p=8.65E-15) (Figure 3.c).

### 2.4 Metastability

We found that the mean metastability was lower in ET (0.1±0.03; p=2.25E-07) compared to control (0.16±0.04). After rTMS stimulation the brain metastability significantly increased (0.18±0.03; p=2.31E-14) (Figure 4.a). These results were supported even at individual subject metastability evaluation (Figure 4.b).

**Figure 4.**
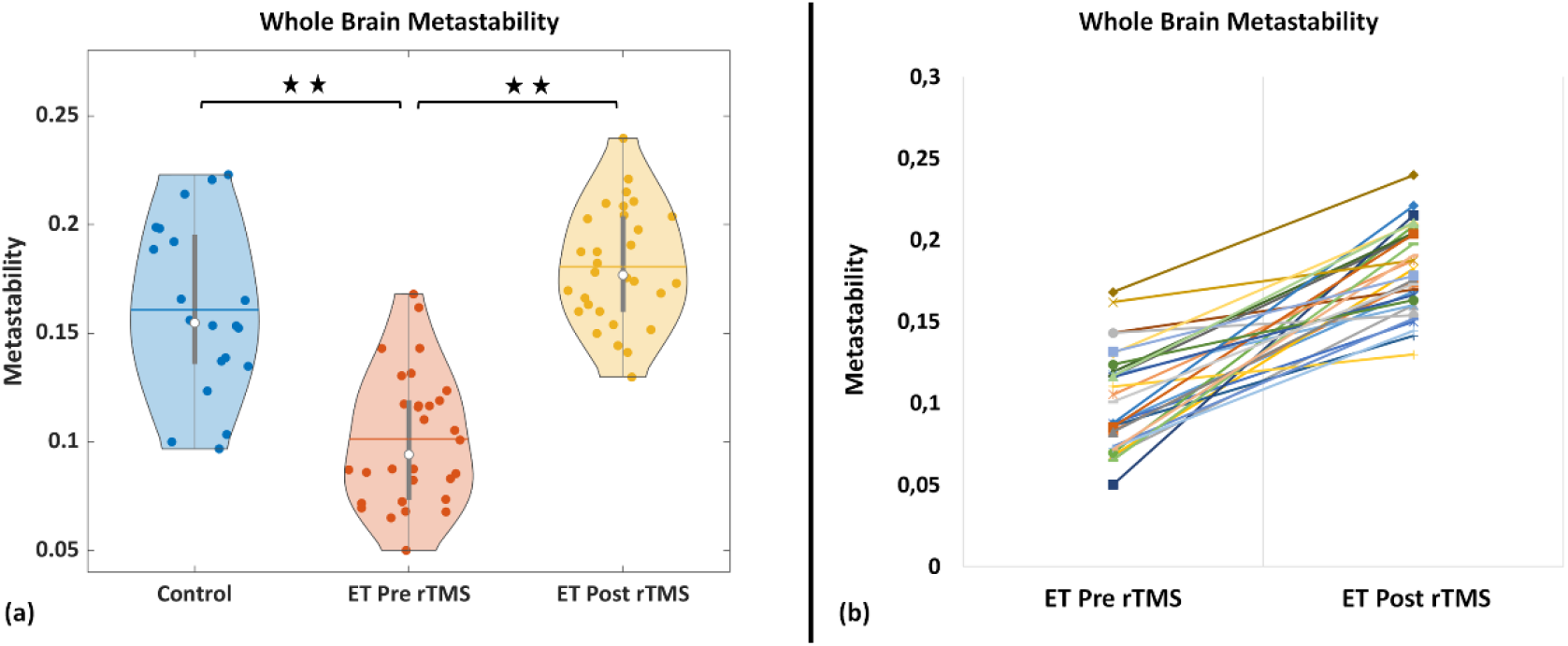
Whole brain metastability (a) in violin plot representations for healthy control, ET pre and post rTMS and (b) individual subject whole-brain metastability pre and post rTMS in ET, depicting significant (* FDR p-value <0.001) increase in metastability after rTMS.

### 2.5 Intrinsic Ignition

We found that whole brain intrinsic ignition was significantly lower in ET (0.8±0.008; p=2.48E-10) when compared to control (0.84±0.02). After rTMS stimulation, it significantly increased (0.84±0.018; p=7.05E-14) (Figure 5.a). We also analyzed region wise intrinsic ignition and noted, several brain regions revealed increased intrinsic ignition after rTMS [Supplementary Figure 1]. Further looking at individual subjects, we noted ET had increased whole brain ignition driven mean integration (IDMI) after rTMS stimulation (Figure 5.b).

**Figure 5.**
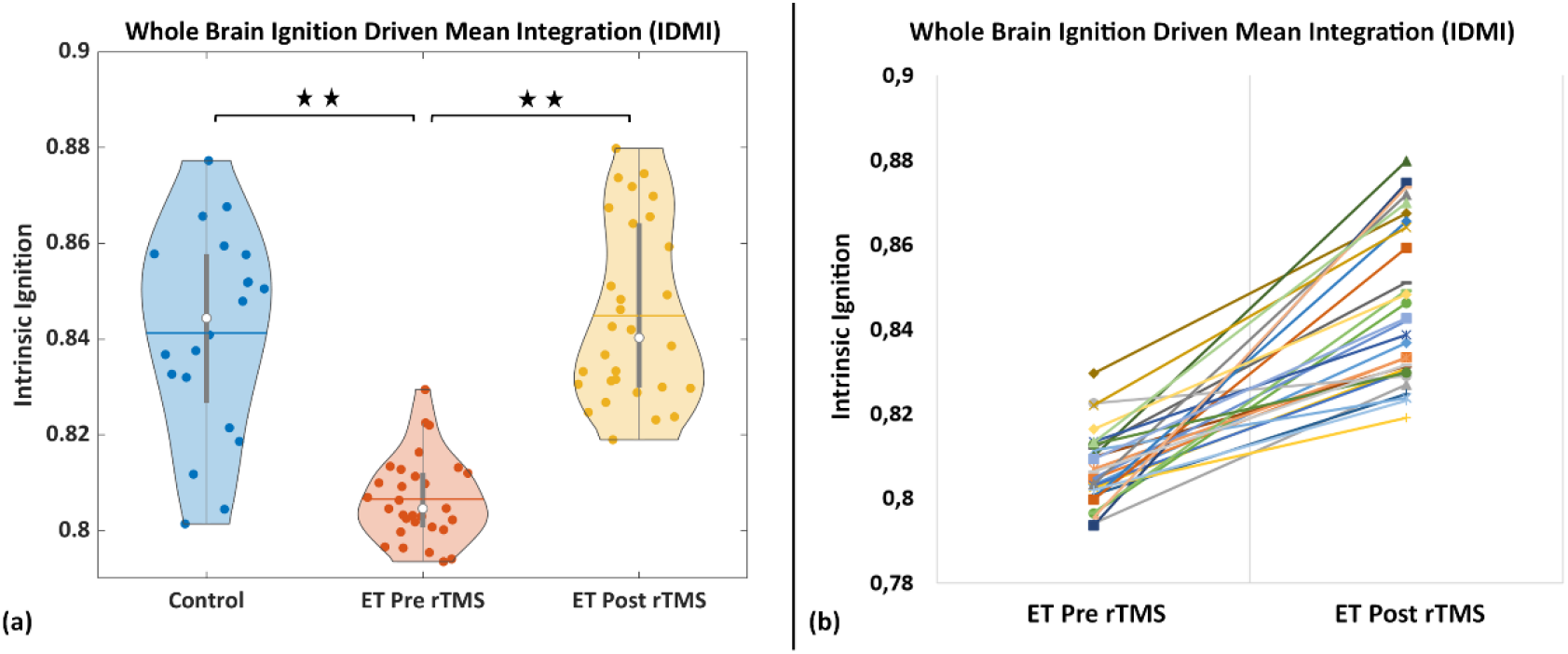
Whole Brain Ignition Driven Mean Integration (IDMI) (a) in violin plot representations for healthy control, ET pre and post rTMS and (b) individual subject whole-brain IDMI pre and post rTMS in ET, depicting significant (* FDR p-value <0.001) increase in IDMI after rTMS.

## 3. Discussion

In this study, we used LF-rTMS in patients with ET to understand its network neurobiology, within the framework of brain network connectivity, integration-segregation, intrinsic ignition, and metastability. In individuals with ET compared to HC at baseline, we saw considerably weaker integration, more segregation, low metastability, and low intrinsic ignition. Unbalanced network integration-segregation, such as weaker integration and excess segregation can result in a less efficient neural communication. Low metastability means that the patterns of connectivity between different brain regions change more frequently and unpredictably, making it more difficult to maintain a consistent pattern of information processing. Low intrinsic ignition means that a brain region is restricted in its capacity to generate and sustain spontaneous neural activity on its own and lacks to propagate those activity to other brain regions ^34,37,39,40^. A single rTMS session was found to reset these metrics by raising global integration, metastability, and intrinsic ignition in ET to levels that were similar to those of healthy controls.

Simulated computational models on metastability reveal small worldness and degree of the structural connectivity to have positive correlation to metastability ^41^. Increased pathlength was associated with a decrease in metastability ^42^. In the current study, ET group revealed reduced integration with reduced metastability compared to controls which synchronously increased after rTMS. These coincident evidences are in line with the evidence from computational models and support increased global integration between nodes as one of the major determinants for improved metastability. Our prior work in patients with Writer’s cramp (WC) using static connectivity methods had revealed increased clustering coefficient, increased small worldness and reduced pathlength after single session of rTMS ^23^ reflecting an increased network integration after rTMS. Not discrediting the obvious differences in analyses methods between static connectivity and DFC we hypothesize that LF-rTMS improves the whole brain network integration, as this feature was consistent across techniques.

Further exploring the spatio-temporal aspects of brain dynamics, we found that rTMS increased brain metastability and intrinsic ignition. Given credence to the paucity of literature in this field using this technique, we looked at the behavioural correlates of such alterations. According to a study on diffuse axonal injury, structural disconnection, decreased cognitive flexibility, and information processing were all associated with altered metastability ^42^. Decreased metastability and ignition has been reported in unresponsive wakeful state and have been found to increase as patients regain consciousness or reach minimal conscious state ^41,43^. Neural responses to perturbations (i.e., TMS) in patients with unresponsive wakeful states, in contrast to patients in minimally conscious states ^44,45^ provides proof of evidence that rTMS increases brain activity and connectivity and supports findings in the current study. Since metastability and intrinsic ignition increased in ET, it is possible that the measure may have the ability to evaluate the impact of rTMS at an individual subject level. We did not record changes in behavioral or cognitive scores after rTMS since clinically evident changes after a single session of rTMS is less known. However, it seems reasonable to assume that increased metastability and intrinsic ignition measures could be further explored to ascertain its comparability with observed improvement in clinical scores in patients undergoing rTMS intervention.

The study had few limitations. First, the effect of sham stimulation in brain dynamics was not evaluated, and all patients were given real LF-rTMS intervention. Second, we did not assess the effect of LF-rTMS stimulation on healthy controls. Third, the described metastability alterations that we have observed are within 10 minutes of a single session of rTMS. It will be interesting to consider changes during stimulation and assess how long these alterations persist to ensure that these changes are not due to anxiety, pain, or fatigue that are associated with rTMS.

## 4. Conclusion

Our findings offer substantial evidence that rTMS altersdynamic functional connectivity measures of metastability and intrinsic ignition. However, further research is necessary to make sure that these alterations are sustainable and can quantify the magnitude of changes required for clinically significant improvement.

## 5. Method

### 5.1 Participants

Thirty patients with ET [mean (age ± SD 37.33) ± 10.67 years, 6 Females] and twenty age, and education-matched HCs [mean (age ± SD) 38.2 ± 10.7 years, 4 Females] participated in this study after providing written informed consent. The study was approved by the institutional ethics committee for humans (NIMHANS, Bengaluru, India). All methods were carried out in compliance with the institutional ethical norms and regulations for human participants. All participants were evaluated in detail by movement disorders specialists [(PKP, RY, NM, and NK); Supplementary Table 1]. All participants were right-handed and were checked for MRI and TMS contraindications. Subjects with ET were either on propranolol, primidone or clonazepam and these drugs were withheld prior to evaluation based on their half-life, i.e., at least 12 hours after the last dose of propranolol or primidone and 40 hours after the last dose of clonazepam. Secondary causes of tremors were primarily ruled out during clinical evaluation, and other causes, for instance hyperthyroidism where suspected was ruled out via blood investigations such as a thyroid profile. Participants with a structural lesion on MRI, prior brain, spinal or peripheral nerve trauma/surgery, claustrophobia and on neuroleptic drugs were excluded from the study. HCs with no neurological or psychiatric illnesses were recruited for MRI.

### 5.2 Experimental Design

Thirty patients of ET, diagnosed as per the consensus criteria of tremor ^46,47^, and twenty, age and gender matched healthy controls were recruited from the neurology outpatient department at NIMHANS. The standard protocol followed for all patients of ET was as follows – informed written consent, clinical evaluation, estimation of resting motor threshold (RMT), resting state functional MRI siting 1 (RsfMRI-s1), rTMS, and a second resting state functional MRI siting 2 (RsfMRI-s2) within 10 minutes of the rTMS. HCs underwent only a single session of resting state functional MRI (RsfMRI) and did not undergo rTMS as ethical approval could not be obtained.

#### 5.2.1 RsfMRI Data Acquisition

A 3T MRI scanner (Skyra; Siemens, Erlangen, Germany) was used for conducting this study. Data was collected between 2016-2019. The acquisition parameters were identical for RsfMRI-s1, RsfMRI-s2 in ET and for RsfMRI in HCs. To prevent head movement, sufficient padding and ear plugs were provided to all subjects. Whole brain Blood oxygen level dependent (BOLD) images were acquired using a spin echo sequence (TR = 2000 ms; TE = 20 ms; refocusing pulse 90°; 4.0 mm slice thickness in an inter-leaved manner with an FOV of 192 × 192 mm^2^; matrix 64 × 64 voxels; voxel size 3 × 3 × 4 mm^3^; 250 TRs). A three-dimensional magnetization-prepared rapid acquisition gradient echo (MPRAGE) sequence was acquired (TR=1900 ms; TE=2.4 ms; voxel size 1×1×1 mm^3^, slice thickness=1mm) for spatial registration and segmentation.

#### 5.2.2 rTMS Parameters

After RsfMRI-s1, subjects were moved to another room adjacent to MRI, and rTMS was delivered using a Magstim Super Rapid stimulator (Magstim Co. Ltd, Whitland, UK) with a figure-of-eight coil configuration. rTMS was applied tangentially to the scalp with the handle pointing backward and laterally at an approximate angle of 45° to the mid-sagittal line, perpendicular to the presumed direction of the central sulcus. rTMS was delivered over the left primary motor cortex (L-M1) by delivering 900 stimuli at 90% of resting motor threshold (RMT) and 1 Hz for 15 min. The RMT was determined as the lowest intensity that produced motor evoked potentials of *>*50 μV in at least five out of 10 consecutive single-pulse TMS stimuli using same Magstim Super Rapid stimulator. The stimulator was attached to an electromyography machine from the right hand first dorsal interosseous muscle using Ag-AgCl surface electrodes placed over the muscle in a belly-tendon arrangement.

### 5.3 Data Analysis

RsfMRI data was recorded for 250 TRs (∼ 8.33 min), however, we removed first 5 TRs from fMRI before pre-processing to avoid MRI scanner signal inhomogeneity due to scanner start transition period.

#### 5.3.1 RsfMRI data pre-processing

RsfMRI data were pre-processed using SPM 12 toolbox (version 7771) with a classical approach of the following steps: realignment, segmentation of the structural data, normalization to MNI152 standard space of 3 × 3 × 3 mm^3^, and head motion (translational and rotational) correction using Friston’s 24-motion parameter ^23^. Data was checked for head motion using the Artefact Detection Toolbox [(ART) (RRID:SCR_005994) (http://web.mit.edu/swg/software.htm)] and found not being significantly different between RsfMRI-s1, RsfMRI-s2 and RsfMRI of HC (for details please refer to Supplementary Table 2).

#### 5.3.2 Brain Region Parcellation

Shen’s 268 region atlas ^48^ was used to parcellate the brain regions into 268 functionally segregated ROIs using MarsBaR toolbox [0.43 Release] ^49^. The RsfMRI BOLD time series for each ROI was extracted by taking average of all the voxels in that ROI.

#### 5.3.3 L-M1 static connectivity

To understand how the L-M1 [Left primary motor cortex area; Broadman area 4; Reference in Shen Atlas: (ROI 158) L.BA4.1] connectivity changes after rTMS induced in the L-M1 brain area (seeds/ROI), we first assessed the static connectivity of L-M1 (i.e., left primary motor cortex area; Broadman area 4) brain regions’ static (i.e., time average) connectivity using seed-to-voxel-based connectivity using functional connectivity toolbox (CONN, http://www.nitrc.org/projects/conn). As per the standard recommendation of CONN, head motion correction was performed for translational and rotational parameters as covariates and the BOLD time series were band-pass filtered with 0.009–0.09 Hz. In addition, to reduce the physiologic noise, CompCor algorithm was used on the segmented white matter and CSF ^50^. BOLD time series in L-M1 seeds was extracted and correlation coefficients between the seed time series and the time series of all other voxels in the brain was derived. Fisher’s r-to-z transformation was used to convert the correlation coefficients to z-scores. Then the general linear model was designed to determine the significant BOLD signal correlation between the mean time series of L-M1 brain seed ROI and that of every other brain voxel at the individual subjects’ level analysis. Second-level random-effects analysis was used to investigate connectivity within conditions (i.e., pre-rTMS vs post-rTMS) and to identify regions with differential connectivity between the two conditions ^51^.

#### 5.3.4 Phase-locking matrices and functional connectivity dynamics (FCD)

To evaluate the level of synchrony in brain activity, we computed the functional connectivity dynamics (FCD) matrices using instantaneous phase synchronisation of the signal at every time point (i.e., dynamics) as described by Deco and colleagues ^41,52^. We first filtered the BOLD time series using a band-pass filter (0.03–0.08 Hz) and demean the BOLD time series prior to extraction of the instantaneous phase *ϕ*_*k*_(*t*) for each BOLD time series (region) ‘k’ ^53^. We computed the instantaneous *ϕ*_*k*_(*t*) using the Hilbert transform ‘*H’* to derive the associated analytical signal of the BOLD time series. The analytical signal represents a narrowband signal, *s*(*t*), in time domain as a rotating vector with an instantaneous phase, *ϕ*(*t*) and an instantaneous amplitude, *A*(*t*), i.e., *s*(*t*) = *A*(*t*). *cos* (*ϕ*(*t*)). The instantaneous phase and the amplitude are given by the argument and the modulus, respectively, of the complex signal, *z*(*t*), given by *z*(*t*) = *s*(*t*) + *i. H*[*s*(*t*)], where *i* is the imaginary unit and *H*[*s*(*t*)] is the Hilbert transform of *s*(*t*). The synchronization between pairs of brain regions was characterized as the cosine of the modulus of difference between their instantaneous phases. For each time-point, the instantaneous phase difference *P*_*jk*_(*t*) between two regions *j* and *k* was calculated as follows:

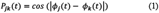

Here, *P*_*jk*_(*t*) = 1 when the two regions are fully synchronized (in phase, *ϕ*_*j*_(*t*) = *ϕ*_*k*_(*t*)), *P*_*jk*_(*t*) = 0 for no synchronization (i.e., orthogonal), and *P*_*jk*_(*t*) = −1 in the antiphase condition^41,52^. P(t) is the phase-interaction matrix at a given time t, which represents the instantaneous phase synchrony among the different brain regions (ROIs). We used these phase interaction connectivity matrices to assess the presence of repeating patterns of network states by computing the recurrence of the phase-interaction patterns called as FCD. The FCD is similar to sliding window and in our study we consider the duration of sliding window to be 30 time points with 1 time point shift ^41,52^. For each time window the average phase-interaction matrix was calculated. Then we constructed the symmetric matrix (MxM), whose (*t*_1_, *t*_2_) entry was defined by the cosine similarity (*S*_cos_) between the upper diagonal elements of phase-interaction matrix ⟨*P*⟩(*t*_1_) *and* ⟨*P*⟩(*t*_2_), as follows:

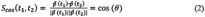

Here, 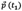 *and* 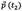 are vectored form of ⟨*P*⟩(*t*_1_) *and* ⟨*P*⟩(*t*_2_) respectively and θ corresponds to the angle formed between the two vectors. The FCD represent the distribution of these cosine similarities for all pairs of time windows.

##### 5.3.4.1 Integration

Integration infers the brain’s capacity to communicate between different parts and subnetworks. We used the phase-locking matrix to compute the level of integration at time *t* based on the procedure presented in ^37^. The integration, *ϕ*, was determined using the largest connected component length of the phase-locking matrix *P*_*jk*_(*t*). More specifically, the phase-locking connectivity matrix was first binarized for a given absolute threshold ‘d’ between 0.01 and 0.99 (0.01≤ d ≤0.99, with an increment of Δd=0.01) and its largest connected component was detected (i.e., the largest sub-group in which any two vertices are connected to each other by paths, and which connects to no additional vertices in the super-graph). The integration, *ϕ*, was defined as the size of the largest connected component ^37,41^.

##### 5.3.4.2 Segregation

Segregation refers to the separation of functional subcomponents (communities/modules) within a system (brain network). Similar to integration, we extracted the community structure of the phase-locking matrix for each time window *t*. Communities were detected using the Louvain algorithm using non-overlapping groups of nodes ^41,54-56^. The modularity index, Q, measures the statistics of the community detection ^55,57^.

##### 5.3.4.3 Metastability

To measure the global level of phase synchronization we used the Kuramoto order parameter ^34,41^, defined as the average phase of the system of N signals (ROIs):

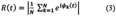

Here, *i* denotes the imaginary unit. For independent signals, the N phases are uniformly distributed and thus R is nearly zero, whereas *R* = 1 if all phases are equal (full synchronization). The metastability measure ^41,52^ quantifies the temporal variability (of this synchronization) *R(t)* and it is measured by its standard deviation.

##### 5.3.4.4 Intrinsic Ignition

Intrinsic ignition describes the influence of spontaneously occurring events within the network over time. The propagation of neural activity was measured using the global integration, (*ϕ*), previously described ^37^, which determines the capacity of the whole network to become interconnected and exchange information. Local events are defined region wise as significantly large fluctuations taking place in the resting-state BOLD signal. To this end, first, the BOLD signals were z-scored (i.e., *Z*_*i*_(*t*)) and then binarized by imposing a threshold θ (i.e., 2 standard deviation above the mean BOLD signal). This resulted in binary time-series per region for which events are indicated with 1 [i.e., *σ*_*i*_(*t*) = 1, if *Z*_*i*_(*t*) > 0 and *σ*_*i*_(*t*) = 0, otherwise ^36^]. Next, for each node ‘*i*’, we calculated *ϕ* in a window of 4 TR after each triggering event (i.e., *σ*_*i*_(*t*) = 1). Finally, the average *ϕ* across all triggering events was calculated to define the Ignition Driven Mean Integration (IDMI) denoted as “Intrinsic Ignition” ^36^.

#### 5.3.5 Statistical Analysis

For the seed-to-voxel static connectivity, voxel-wise paired sample t-test was performed to detect regions with significant differences in connectivity between the two conditions. The statistical threshold was set to a cluster-level-corrected α value of 0.05 for voxel-wise p-value of <0.05 with false-discovery rate (FDR) correction (with a minimum cluster extent of 20 contiguous voxels) ^51^. For the whole brain measures (i.e., mean FCD, Integration, segregation, metastability, and Ignition) we first assessed the normal distribution and noted all these measures were following a normal distribution. Then, to assess the group difference between healthy controls and ET for the above measures, we used a two-tailed, two-sample *t*-test (i.e., ‘ttest2’ of MATLAB function) with p<0.05. For between condition assessment (ET patients pre-rTMS vs post-rTMS), we used a two-tailed, paired sample *t*-test (i.e., ‘ttest’ of MATLAB function) with p< 0.05.

## Acknowledgements

This work was supported by the Department of Science and Technology - Cognitive Science Research Initiative (DST-CSRI), Government of India [Grant number: SR/CSRI/162/2013]. Mr. Sujas Bhardwaj is currently a Senior Research Fellow under the Wellcome DBT India Alliance Intermediate Fellowship [IA/CPHI/17/1/503348] of Dr. Shantala Hegde (Additional Professor, Music Cognition Lab, NIMHANS, Bengaluru-29, India).

## Author Contribution

The experiment was devised and planned by RDB and PKP. The clinical evaluation of the patients’ scores for inclusion criteria was carried out by PKP, RY, NK, AS, and SP. The subjects received the rTMS from AS and SB. The MRI data were collected by SB, AS, SKK, and RDB. VT and KK conducted neuropsychological evaluations. The data was examined by SB, RP, and RDB, and the findings were then explained by RDB, SB, RP, and PKP. The method development was supervised by RDB, PKP, GD, JA, and SL. A critical evaluation was offered by JA, SL, AS, KU, NM, SP, RK, and GD. RDB, SB, and RP wrote the manuscript. The results were interpreted, the manuscript was modified, and it was approved by all authors. None of the authors of this work has any competing interests. SB, RP, and RDB have equally contributed to the manuscript.

## Data and Codes Availability Statement

The data that supports the findings of this study will be available from the corresponding author, [RDB], upon reasonable request and for research purpose only. Please find the relevant codes for the computation of DFC matrices at https://github.com/RajanikantPanda/rTMS_ET_Brain_Metastability_Ignition.

