## Supplementary Material for "Single session of rTMS enhances brain metastability and intrinsic ignition"

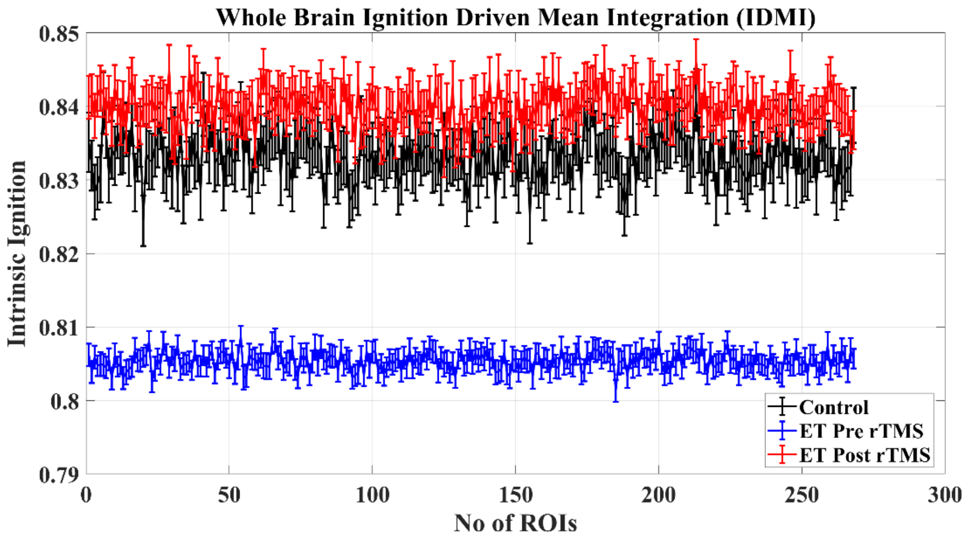


**Supplementary Figure 1.** The region-wise plot of Ignition Driven Mean Integration (IDMI) for healthy controls, ET pre and post-rTMS.

**Supplementary Table 1:** Clinical and cortical excitability measures in subjects with essential tremor

| NAME | Age | Gender | AAO | DD | RMT | AMT | CMCT [ms] | CSP |
|---|---|---|---|---|---|---|---|---|
| Subject 001 | 47 | M | 38 | 9 | 36 | 30 | 7.57 | 98.6 |
| Subject 002 | 24 | M | 20 | 4 | 35 | 31 | 8.6 | 64.8 |
| Subject 003 | 27 | M | 19 | 8 | 42 | 39 | 7.2 | 66.4 |
| Subject 004 | 25 | M | 21 | 4 | 44 | 41 | 9.5 | 52.8 |
| Subject 005 | 36 | M | 26 | 10 | 39 | 32 | 7.8 | 59.3 |
| Subject 006 | 22 | M | 7 | 15 | 38 | 31 | 8.1 | 66.4 |
| Subject 007 | 44 | F | 43 | 1 | 47 | 36 | 8.95 | 67.2 |
| Subject 008 | 32 | M | 17 | 15 | 40 | 34 | 8.2 | 48.4 |
| Subject 009 | 32 | F | 31 | 1 | 38 | 34 | 6.8 | 62.6 |
| Subject 010 | 41 | F | 38 | 3 | 39 | 29 | 6.9 | 67.9 |
| Subject 011 | 50 | M | 36 | 14 | 34 | 30 | 4.7 | 52.8 |
| Subject 012 | 42 | M | 37 | 5 | 30 | 28 | 5.8 | 58.4 |
| Subject 013 | 45 | F | 44.5 | 0.5 | 38 | 29 | 6.6 | 55.3 |
| Subject 014 | 35 | M | 22 | 13 | 39 | 31 | 5.9 | 59.1 |
| Subject 015 | 41 | M | 34 | 7 | 32 | 26 | 7.4 | 42.8 |
| Subject 016 | 28 | M | 13 | 15 | 34 | 30 | 8.45 | 46.4 |
| Subject 017 | 47 | M | 46 | 1 | 70 | 62 | 15.5 | 46.6 |
| Subject 018 | 23 | M | 16 | 7 | 24 | 22 | 7.9 | 37.6 |
| Subject 019 | 32 | M | 28 | 4 | 36 | 32 | 7.5 | 78.4 |
| Subject 020 | 25 | M | 24 | 1 | 32 | 26 | 8.24 | 48.4 |
| Subject 021 | 35 | M | 21 | 14 | 40 | 26 | 6.6 | 117.2 |
| Subject 022 | 55 | M | 54 | 1 | 38 | 36 | 5.4 | 48 |
| Subject 023 | 54 | M | 53 | 1 | 40 | 38 | 6.8 | 86.2 |
| Subject 024 | 20 | F | 14 | 6 | 28 | 25 | 7.4 | 64 |
| Subject 025 | 46 | M | 41 | 5 | 30 | 22 | 9.1 | 52.2 |
| Subject 026 | 33 | M | 31 | 2 | 38 | 32 | 8.3 | 56.7 |
| Subject 027 | 26 | F | 22.5 | 3.5 | 38 | 34 | 8.6 | 46.8 |
| Subject 028 | 49 | M | 44 | 5 | 42 | 32 | 6.8 | 91.5 |
| Subject 029 | 54 | M | 53 | 1 | 44 | 38 | 4.4 | 54 |
| Subject 030 | 51 | M | 49 | 2 | 42 | 34 | 5.4 | 78.6 |

AAO: age at onset; DD: Disease duration; RMT: Resting Motor Threshold, AMT: Active Motor Threshold, CMCT: Central Motor Conduction Time; CSP: Contralateral Silent Period.

**Supplementary Table 2:** Demographic and Head motion values:

| | | HC | ET-PRE | ET-POST | p-value |
|---|---|---|---|---|---|
| **N** | | 20 (4 Females) | 30 (6 Females) | | - |
| **Age (Mean ± STDEV) Years** | | 38.2 ± 10.71 | 37.33 ± 10.67 | | HC vs ET: **NS** |
| **Head Motion** | **Translational (mm)** | x=0.47±0.42; y=0.56±0.33; z=1.17±0.71 | x=0.60±0.48, y=0.62±0.40, z=1.15±0.56, | x=0.69±0.58, y=0.78±0.59, z=1.46±1.08, | HC vs ET-PRE: **NS** p= 0.30; 0.58; 0.90<br>ET-PRE vs ET-POST: **NS** p= 0.11; 0.19; 0.17 |
| | **Rotational (degree)** | pitch=0.022±0.007; roll=0.009±0.007; yaw=0.009±0.007 | pitch=0.020±0.010, roll=0.010±0.006, yaw=0.010±0.011, | pitch=0.024±0.018, roll=0.011±0.010, yaw=0.013±0.01 | HC vs ET-PRE: **NS** p= 0.74; 0.81; 0.77<br>ET-PRE vs ET-POST: **NS** p= 0.19; 0.12; 0.14 |
